# *In-vitro* and *in-vivo* efficacy of a novel broad spectrum β-lactamase inhibitor APC24-7 against *Enterobacterales*

**DOI:** 10.64898/2026.03.04.709740

**Authors:** Carina Matias, Klaus Skovbo Jensen, Bjørg Bolstad, Bjørn Klem, Pål Rongved, Carina Vingsbo Lundberg, Jon Ulf Hansen

## Abstract

The rise of multidrug-resistant (MDR) bacteria, particularly carbapenem-resistant *Enterobacterales* (CRE), poses a significant threat to public health. Infections caused by CRE, such as *Escherichia coli* and *Klebsiella pneumoniae*, are associated with high rates of antibiotic treatment failure. β-lactam antibiotics, like meropenem, remain crucial in treating these infections, but their efficacy is undermined by β-lactamase production. This study investigates the potential of APC24-7, a novel broad-spectrum β-lactamase inhibitor (BLi) with dual activity, to restore antimicrobial activity of meropenem against CRE clinical isolates. The *in-vitro* analysis of a diverse panel of clinically relevant *E. coli* and *K. pneumoniae* isolates expressing both serine- and metallo-β-lactamases demonstrated that APC24-7 effectively restored meropenem activity by reducing the minimum inhibitory concentrations (MICs) to below breakpoint. Time-kill assays confirmed that the combination therapy showed dose-dependent bacterial killing, with significant potentiation of meropenem activity against isolates expressing both serine- and metallo-β-lactamases. *In-vivo* efficacy evaluation in a murine thigh infection model further confirmed APC24-7’s potential to restore meropenem efficacy against meropenem resistant strains. These findings suggest that APC24-7offers a promising strategy to combat infections caused by β-lactamase-producing *Enterobacterales*.

## Introduction

The emergence and worldwide spread of multidrug-resistant bacteria contribute to the leading public health threats of the 21^st^ century. In particular, infections caused by carbapenem-resistant Enterobacterales (CRE), classified by the World Health Organization (WHO) as critical group of priority pathogens, including *Escherichia coli* and *Klebsiella pneumoniae*, have been associated with high rates of antibiotic treatment failure and mortality [1]–[3]. β-lactam antibiotics (BLs), including carbapenems, remain among the most widely used agents in hospital settings due to their efficacy and broad activity spectrum [1], [2]. However, increasing resistance rates threaten their continued effectiveness and highlight the need to develop novel treatment strategies [4], [5]. Resistance to BLs arises by multiple mechanisms, with one of the most clinically relevant being the production of β-lactamases. These enzymes can be classified either in to zinc-ion-dependent metallo-β-lactamases (MBLs, Ambler Class B) or nucleophilic serine-β-lactamases (SBLs, Amber Class A, C and D) [6]–[8], where enzymes belonging to classes A, B and D are frequently associated to carbapenem resistance [9]. Both groups catalyze hydrolysis of the β-lactam ring via distinct mechanisms and differ in their substrate specificity and distribution among bacterial species [7], [10]. Multiple β-lactamases can coexist alongside additional resistance mechanisms and consequently CRE are frequently resistant to multiple antimicrobial classes which significantly limits treatment options [11]–[13].

Significant advancements have been made in the development of novel β-lactamase inhibitors (*BLis*) [4], [14], [15]. These inhibitors are designed to be utilized alongside existing BLs, aiming to protect BLs from degradation and thus, to improve treatment outcomes and mitigate the repercussions of drug-resistant infections [16], [17]. Since 2015 six novel BL/BLi combinations, aztreonam-avibactam [18], [19], ceftadizime-avibactam (CZA), meropenem-vaborbactam, imipenem-relobactam (IMR) [20], sulbactam/durlobactam [21] and cefepime-enmetazobactam, have been approved, providing new therapeutic options for infections caused by CRE. However, these inhibitors primarily target SBL-mediated resistance, particularly classes A and C, with limited activity against Class D enzymes [15].

Infections caused by MBL-producing pathogens remain difficult to treat [15]. Current options are limited to aztreonam combined with avibactam, ceftazidime-avibactam or monotherapy with the novel siderophore cephalosporin cefiderocol [22]–[25]. Though cefiderocol effectively target MBL-producing organisms, resistance has already been reported [26]–[29]

To date, no MBL inhibitors have been approved; however several promising MBL inhibitors have been reported at different stages of development [4], [14], [15]. Compounds such as ANT2681 [30], and indole carboxylate inhibitors [31], have been described. Other BL/*BLi*, combinations including cefepime/taniborbactam [32], cefepime/zidebactam (WCK 5222) [33], cefepime/nacubactam [34], β-lactam/xeruborbactam (QPX7728) [35] or β-lactam/APC148 [36] are undergoing clinical evaluation with some concluding clinical Phase 3 [15], [37].

APC24-7 has been recently reported as a unique molecule that incorporates two inhibitory moieties with complementary modes of action, resulting in a hybrid broad-spectrum inhibitor active against both MBLs and SBLs. APC24-7 can thereby restore the activity of meropenem against bacteria expressing both MBL and SBL enzymes [38].

Here, we describe expanded *in-vitro* microbiological profiling of APC24-7 in combination with meropenem, against *E. coli* and *K. pneumoniae* clinical isolates harboring multiple MBL and/or SBL enzymes and provide *in-vivo* efficacy proof-of-concept in a murine thigh infection model.

## Results

### APC24-7 restores susceptibility to meropenem

The MIC values of meropenem and APC24-7 were determined for a panel of fifty clinical and reference *K. pneumoniae* and *E. coli* isolates producing a broad range of MBL and/or SBL. All isolates were classified as resistant to meropenem with MIC ≥ 4 mg/L. No intrinsic antibacterial activity of APC24-7 was observed at concentrations up to 64 mg/L (data not shown). To assess the ability of APC24-7 to restore meropenem activity, a checkerboard assay was performed using increasing APC24-7 concentrations in combination with meropenem against 31 *K. pneumoniae* strains (Table 1) and 17 *E. coli* strains (Table 2). APC24-7 restoration of meropenem activity was dose dependent with several strains becoming susceptible (CLSI susceptibility breakpoint, MIC ≤ 1 mg/L) [39] with APC24-7 concentrations as low as 4–8 mg/L. Figure 1 summarizes the checkerboard assay results with the APC24-7 concentration of 32 mg/L showing restored susceptibility to meropenem for 11 out of 17 *E. coli* isolates including isolates harboring OXA-48 or NDM-5 carbapenemases. For *K. pneumoniae* 16 out of 31 clinical isolates remained resistant to meropenem.

**Table 1.** *K.pneumoniae* clinical isolates meropenem susceptibility profile, monotherapy and in combination with APC24-7.

| Isolate ID | β-lactamase(s) | Meropenem MIC (mg/L) |  |  |  |  |  |
| --- | --- | --- | --- | --- | --- | --- | --- |
|  |  | - | APC24-7 (mg/L) |  |  |  |  |
|  |  |  | 4 | 8 | 16 | 32 | 64 |
| 76 | VIM-1, SHV-30 | 32 | 0.25 | 0.25 | ≤ 0.06 | ≤ 0.06 | ≤ 0.06 |
| AMA2650 | KPC-3, SHV-62, TEM-214, TEM-213, TEM-210, TEM-209, TEM-208, TEM-206, TEM-141, TEM-57, TEM-34, TEM-33 TEM-1B, OXA-9, CTX-M-15 | 128 | 0.5 | 0.5 | 0.25 | 0.25 | 0.125 |
| DIH | VIM-19, SHV-172, TEM-1B, CMY-4, CTX-M-3 | 64 | 1 | 0.5 | 0.125 | ≤ 0.06 | ≤ 0.06 |
| AMA2119 | KPC-3, CTX-M-15, OXA-1, OXA-9, SHV-28, SHV-106, TEM-1B, TEM-33, TEM-34, TEM-141, TEM-206, TEM-209, TEM-210, TEM-214, TEM-216 | 128 | 4 | 2 | 0.5 | 0.125 | 0.125 |
| 135 | VIM-1, SHV-12, TEM-1A, OXA-9 | 32 | 4 | 2 | ≤ 0.06 | ≤ 0.06 | ≤ 0.06 |
| 160 | OXA-48, SHV-11 | 16 | 4 | 2 | ≤ 0.06 | ≤ 0.06 | ≤ 0.06 |
| ATCCBAA1705 | KPC-2 | 64 | 4 | 2 | 0.5 | 0.25 | 0.125 |
| 34 | IMP-4, SHV-11, TEM-1 | 16 | 8 | 8 | 8 | 2 | ≤ 0.06 |
| AMA004435 | OXA-48, SHV-26-like | 128 | 16 | 4 | 1 | 0.5 | 0.125 |
| B68-1 | NDM-1, AmpH, CTX-M-15, CMY, OXA-9, SHV-33, TEM-168 | 128 | 16 | 4 | 1 | ≤ 0.06 | ≤ 0.06 |
| ST258 | KPC-2, OXA-9 like, SHV-5-like, TEM-1A-like | 64 | 16 | 16 | 2 | 0.5 | 0.125 |
| SECURE ST147 | NDM-1, KPC-2 | 64 | 16 | 8 | 4 | 0.5 | 0.25 |
| AMA877 | KPC-3, OXA-1, SHV-182, SHV-159, SHV-158, OXA-9 | 128 | 16 | 16 | 4 | 0.25 | 0.25 |
| 80 | IMP-4, OKP-B-21, OXA-1, SFO-1 TEM-1 | 16 | 16 | 16 | 16 | 4 | 0.125 |
| AMA004417 | VIM-1, KPC-2, SHV-99, TEM-1B, OXA-10 | 1024 | 32 | 16 | 8 | 0.5 | 0.125 |
| 850 | OXA-1, OXA-48, SHV-11, TEM-1 | 64 -128 | 32 | 32 | 32 | 32 | 32 |
| 1149 | IMP-18, KPC-45, OXA-1, OXA-2, SHV-1, CTX-M-15 | 32 | 32 | 32 | 32 | 32 | ≤ 0.06 |
| AMA004817 | NDM-9, OXA-1, TEM-1B, SHV-28, SHV-106, CTX-M-15 | 128 | > 64 | > 64 | 16 | 0.5 | ≤ 0.06 |
| NCTC13443 | NDM-1, CMY-4, CTX-M-15, SHV-106, SHV-28, OXA-1, DHA-24, DHA-1, TEM1-B, TEM1-A, SHV-1A, SHV-185, SHV-11, SHV-70, SHV-69, SHV-31, SHV-25, SHV-13, OXA-9 | 512 | > 64 | > 64 | > 64 | 4 | 1 |
| 40 | VIM-27, CTX-M-15, OXA-1, SHV-11 | 1024 | > 64 | > 64 | > 64 | 32 | 1 |
| AMA003821 | KPC-2, SHV-12 | 1024 | > 64 | > 64 | > 64 | 64 | 16 |
| AMA1348 | KPC-2, OXA-9, TEM-79, SHV-182, SHV-158 | 1024 | > 64 | > 64 | 64 | 16 | 8 |
| CPO20170084 | NDM-1, OXA-48, OXA-9, SHV-106, SHV-28, TEM-1A, CTX-M-15 | 256 | > 64 | > 64 | 64 | 32 | 16 |
| AMA004208 | NDM-4, OXA-181, TEM-1B, SHV-199, SHV-194, SHV-179, SHV-145, SHV-98, SHV-78, SHV-26 | 1024 | > 64 | > 64 | > 64 | 32 | 8 |
| AMA004950 | NDM-5<br>OXA-232, SHV-182-like, CTX-M-15, TEM-1B | >1024 | > 64 | > 64 | > 64 | 64 | 32 |
| AMA004443 | NDM-1, KPC-2, CTX-M-15, TEM-1B, TEM, 30, TEM-104, TEM-198, TEM-207, TEM-217, TEM-230, TEM-34 | 1024 | > 64 | > 64 | > 64 | 8 | 2 |
| AMA1498 | KPC-3, SHV-28-like | 1024 | > 64 | > 64 | 64 | 16 | 8 |
| 1158 | NDM-5, VIM-2, CTX-M-15, OXA-1, OXA-232, SHV-11, TEM-1 | 512 | > 64 | > 64 | > 64 | 64 | 16 |
| ST147 | OXA-48, CTX-M-15, VEB-8, SHV-11, SHV-67, CMY-2, OXA-1, TEM-1B | 64 | 64 | 64 | 64 | 64 | 64 |
| AMA005047 | NDM-5, OXA-9, OXA-48, SHV-11, SHV-67, TEM-1C, CTX-M-15, CTX-M-14b | 256 | > 64 | > 64 | > 64 | > 64 | > 64 |
| 556 | CTX-M-15, OXA-48, OXA-9, SHV-100, TEM-1A | 16-32 | 16 | 16 | 16 | 16 | 16 |

**Table 2.** *E.coli* clinical isolates meropenem susceptibility profile, monotherapy and in combination with APC24-7.

| Isolate ID | β-lactamase(s) | Meropenem MIC (mg/L) |  |  |  |  |  |
| --- | --- | --- | --- | --- | --- | --- | --- |
|  |  | - | APC24-7 (mg/L) |  |  |  |  |
|  |  |  | 4 | 8 | 16 | 32 | 64 |
| AMA004326 | KPC-3, OXA-9-like, TEM-1A-like | 32 | 0.125 | 0.125 | 0.125 | 0.125 | 0.06 |
| AMA003244 | NDM-5, OXA-232, CTX-M-15, TEM-1B | 64 | 1 | 0.5 | 0.5 | 0.5 | 0.125 |
| 149 | NDM-7, CMY-42 | 64 | 4 | 2 | 2 | 2 | 1 |
| AMA817 | NDM-1, CMY-6, DHA-1, OXA-1, TEM-1-C | 32 | 8 | 2 | 1 | 0.125 | 0.06 |
| AMA005000 | NDM-5, OXA-48, CTX-M-3, TEM-1B | 128 | 8 | 4 | 2 | 1 | 0.5 |
| AMA004995 | NDM-5, OXA-181, CMY-42 | 256 | 8 | 8 | 2 | 1 | 0.25 |
| 1162 | NDM-5, CTX-M-15, OXA-48 | 256 | 16 | 8 | 8 | 2 | 0.5 |
| AMA005098 | NDM-5, KPC-3, SHV-182-like, CTX-M-15, CMY-145, TEM-1B | 256 | 32 | 8 | 4 | 0.5 | 0.25 |
| IR57 | NDM-1, CTX-M-15, TEM-198, AmpC, OXA-1, FonA-2 | 256 | 32 | 16 | 2 | 1 | 0.125 |
| IHMA997800 | NDM-5, CTX-M-15, TEM-1-B, CMY-2 | 256 | 32 | 32 | 8 | 4 | 2 |
| AMA003170 | NDM-5, OXA-181, CTX-M-15, CMY-2, OXA-1, TEM-1B | 32 | 32 | 16 | 8 | 4 | 1 |
| CPO20180141 | NDM-5, OXA-244, CTX-M-15, OXA-1 | 32 | 32 | 8 | 4 | 1 | 0.125 |
| AMA005043 | NDM-5, OXA-244 | 128 | 64 | 64 | 16 | 4 | 0.25 |
| Espagne | VIM-1, TEM-1B | 32 | 64 | 64 | 16 | 1 | 0.25 |
| 162 | NDM-7, CTX-M-15, TEM-1B | 128 | > 64 | 64 | 32 | 8 | 0.125 |
| OM20 | NDM-1, CTX-M-15, TEM-1 | 128 | 128 | 64 | 16 | 1 | 0.25 |
| IHMA997577 | NDM-5, CTX-M-15, TEM-1-B | 256 | 256 | 64 | 32 | 2 | 0.25 |
| ATCC 25922 | - | 0.03 | -- | -- | -- | -- | -- |

**Interpretation Tables 1 & 2:** Grey: no changes in MIC were observed; Green: susceptible, MIC below susceptibility breakpoint ( $\leq 1$ mg/L); Yellow: intermediate susceptibility ( $=2$ mg/L); Red: resistant, MIC above resistance breakpoint ( $\geq 4$ mg/L).

**Figure 1.**
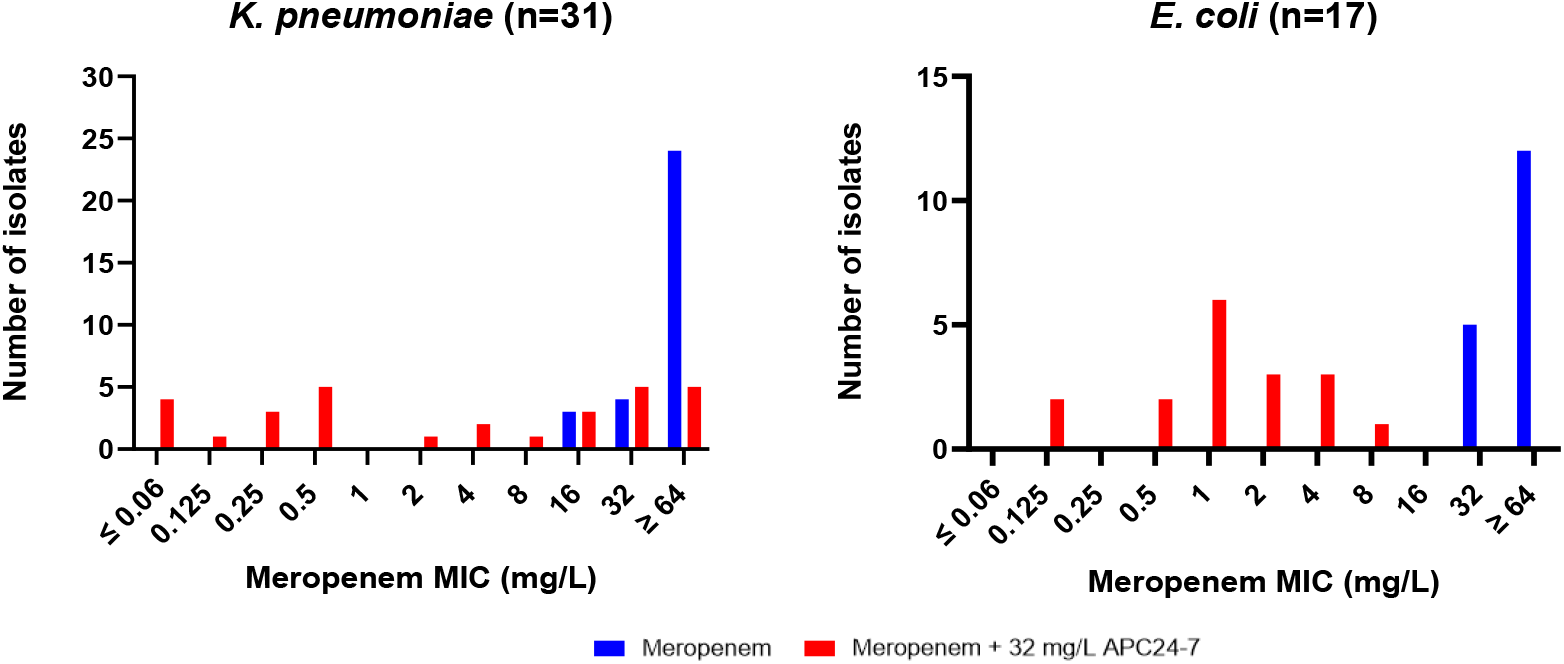
Meropenem MIC distribution, both monotherapy and in combination with 32 mg/L APC24-7 in *K. pneumoniae* (n=31) and *E. coli* (n=17). Blue bars indicate Meropenem alone; red bars indicate Meropenem combined with 32 mg/L APC24-7. A marked reduction in meropenem MIC is overall observed, demonstrating the potentiating effect of APC24-7.

### Kill-kinetics of meropenem in combination with APC24-7

To further evaluate the growth inhibition of the combination of meropenem and APC24-7, time-kill studies were conducted using four *E. coli* and five *K. pneumoniae* meropenem resistant isolates (MIC ≥ 32 mg/L) identified in the checkerboard assay, as being sensitive to meropenem in combination with 64 mg/L APC24-7 or lower and carrying an MBL and/or SBLs of clinical relevance (including NDM, VIM, IMP, OXA, KPC). Bacterial cultures were exposed to varying concentrations of meropenem combined with a fixed concentration of 32 mg/L APC24-7 over a 24-hour period. In combination with APC24-7 at 32 mg/L, meropenem resulted in a bactericidal activity, defined as a decrease of ≥ 3 log_10_ CFU/mL compared to the initial bacterial count [40]. This bactericidal activity was observed within 3 to 5 hours for all 9 selected clinical isolates (Figure 2) For the 5 *K. pneumoniae* isolates, 1 mg/L of meropenem was sufficient for bactericidal activity in combination with 32 mg/L APC24-7 whereas for some of the *E. coli* isolates 4-16 mg/L of meropenem was required.

**Figure 2.**
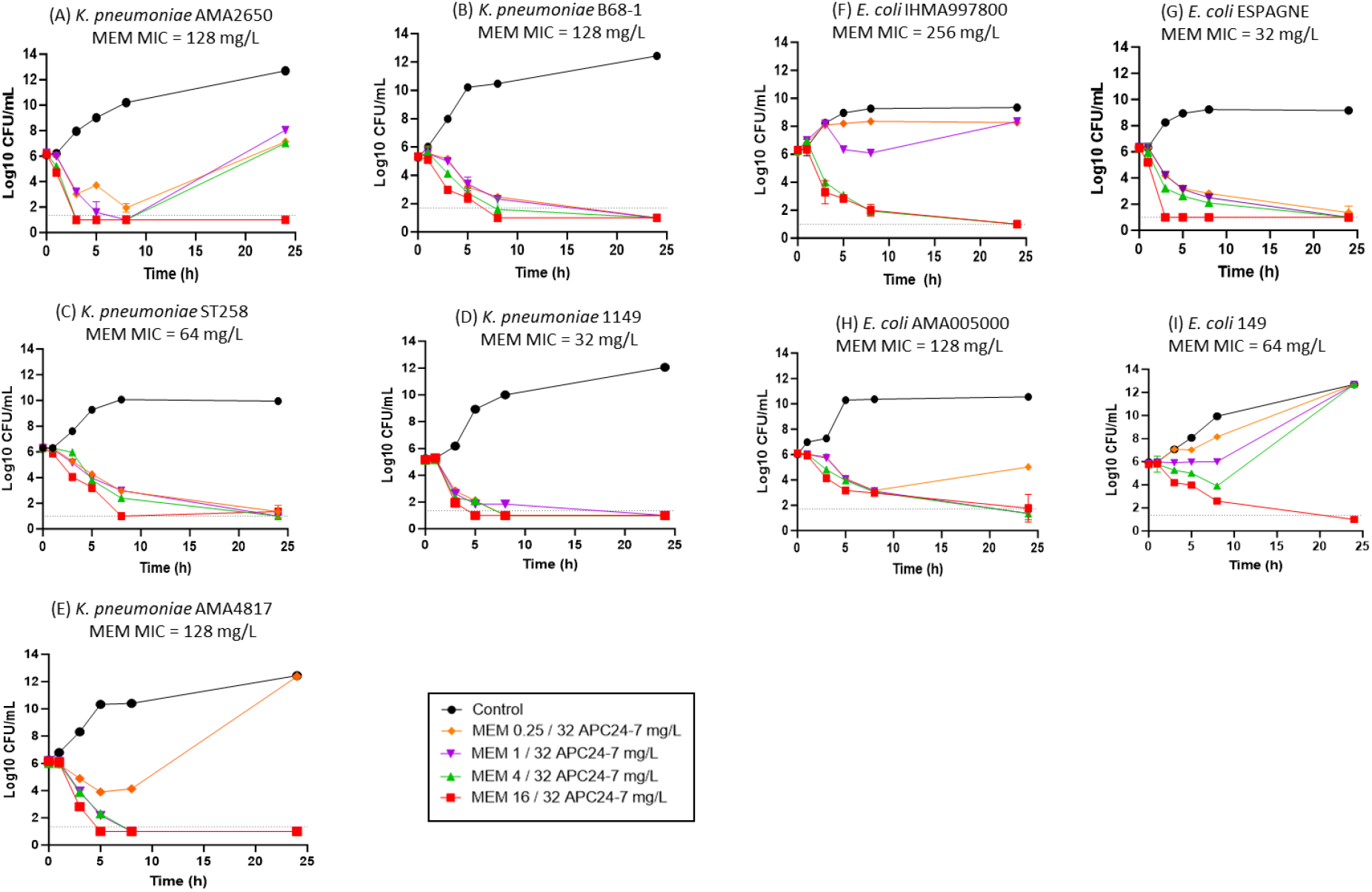
Time-kill curves of meropenem resistant clinical isolates (MIC ≥ 32 mg/L) (A) *K. pneumoniae* AMA2650, (B) *K. pneumoniae* B68-1, (C) *K. pneumoniae* ST258, (D) *K. pneumoniae* 1149 and (E) *K. pneumoniae* AMA4817 (F) *E. coli* IHMA997800, (G) *E. coli* Espagne, (H) *E. coli* AMA005000, (I) *E. coli* 149, treated with meropenem at 0.25, 1, 4 and 16 mg/L in combination with a fixed concentration of APC24-7 (32 mg/L). Growth controls (drug free media) were included in all experiments. Dotted line on each graph indicates the lower detection limit. Data is shown as the average +/- SD log10 CFU/mL of duplicates per time point.

### Killing rate of meropenem in combination with APC24-7

To further investigate the effect of APC24-7 and meropenem on bacterial growth, a two-dimensional scheme was applied to four sets of time-kill curves for *E. coli* IHMA997800 (Figure 3). This way, an estimate could be obtained for the instantaneous growth rate as a function of the static, simultaneous concentrations of meropenem and APC24-7, as shown in Figure 4. For the initial net growth rate *μ* of *E. coli* IHMA997800, the concentration of APC24-7 has a pronounced influence on the meropenem EC50 and the apparent sigmoid effect curve. The pharmacodynamic MIC, *zMIC*, (i.e., the meropenem concentration at which *μ* = 0), is closely related to EC50 and decreases from 4.3 mg/L to 0.35 mg/L when the APC24-7 concentration goes from 2 mg/L to 128 mg/L (Figure 4A). If a 5-hour initial time period is used (instead of 3 hours) for the estimation of *μ*, the fitted sigmoid curves show a decrease in the *zMIC* from 3.4 mg/L to 0.32 mg/L for the above-mentioned shift in APC24-7 concentration (Figure 4B).

**Figure 3.**
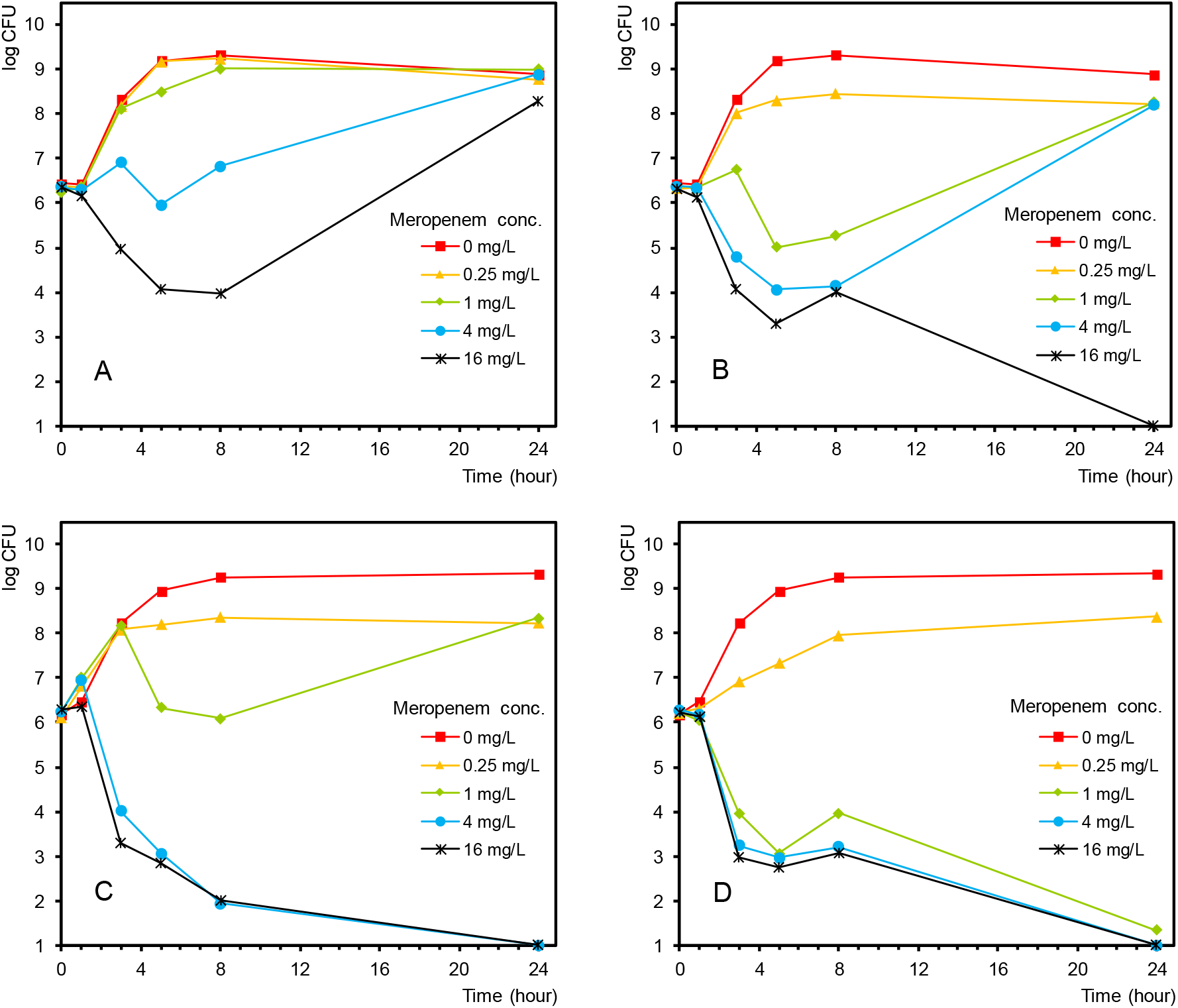
Time-kill curves of *E. coli* IHMA997800 treated with meropenem at 0.25, 1, 4 and 16 mg/L in combination with APC24-7 at (A) 2, (B) 8, (C) 32 and (D) 128 mg/L.

**Figure 4.**
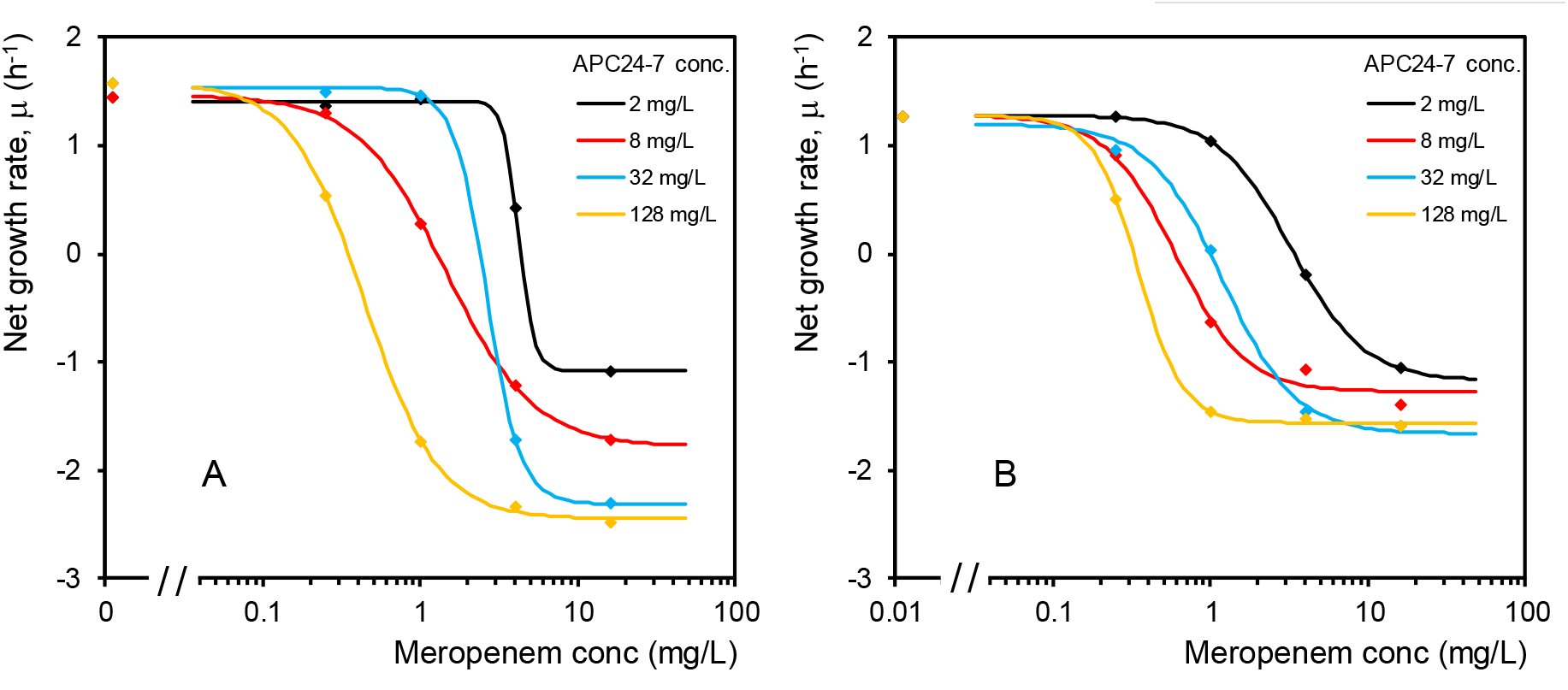
Initial net growth rate of *E. coli* IHMA997800 for various combinations of meropenem and APC24-7 concentrations. The net growth rate is shown as its mean value in the period t = 0 - 3h (A) and t = 0 - 5h (B) of the time-kill experiments.

### APC24-7 pharmacokinetics

Pharmacokinetic (PK) profiles of three dose levels of APC24-7 (3, 10, 30 mg/kg) given subcutaneous (SC) were generated in neutropenic thigh-infected female mice (Figure 5). None of the two PK parameters, C_max_ or AUC_inf_, were proportional to dose, with higher doses indicating slower and/or incomplete absorption (Table 3). Plasma levels of ≥16 mg/L APC24-7, was achieved only with the higher dose of 30 mg/kg, resulting in sustained plasma levels >20 mg/kg for more than an hour after dose (Figure 5). Relative to meropenem, APC24-7 had a long terminal half-life of 40-50 minutes (Table 3).

**Table 3.** Plasma PK parameters from noncompartmental analysis of APC24-7 in thigh-infected female NMRI mice given a SC dose.

| APC24-7 dose | T<sub>1/2</sub> (h) | T<sub>max</sub> [h] | C<sub>max</sub> [mg/L] | AUC<sub>inf</sub> [h·mg/L] | V<sub>ss/w</sub> [L/kg] | CL/w [L/(h·kg)] | kel' [1/h] |
| --- | --- | --- | --- | --- | --- | --- | --- |
| 30 mg/kg SC | 0.77 | 1.0 | 26 | 51 | 0.80 | 0.58 | 0.73 |
| 10 mg/kg SC | 0.79 | 0.25 | 14 | 19 | 0.65 | 0.53 | 0.82 |
| 3 mg/kg SC | 0.66 | 0.25 | 7.7 | 8.5 | 0.37 | 0.35 | 0.95 |

**Figure 5.**
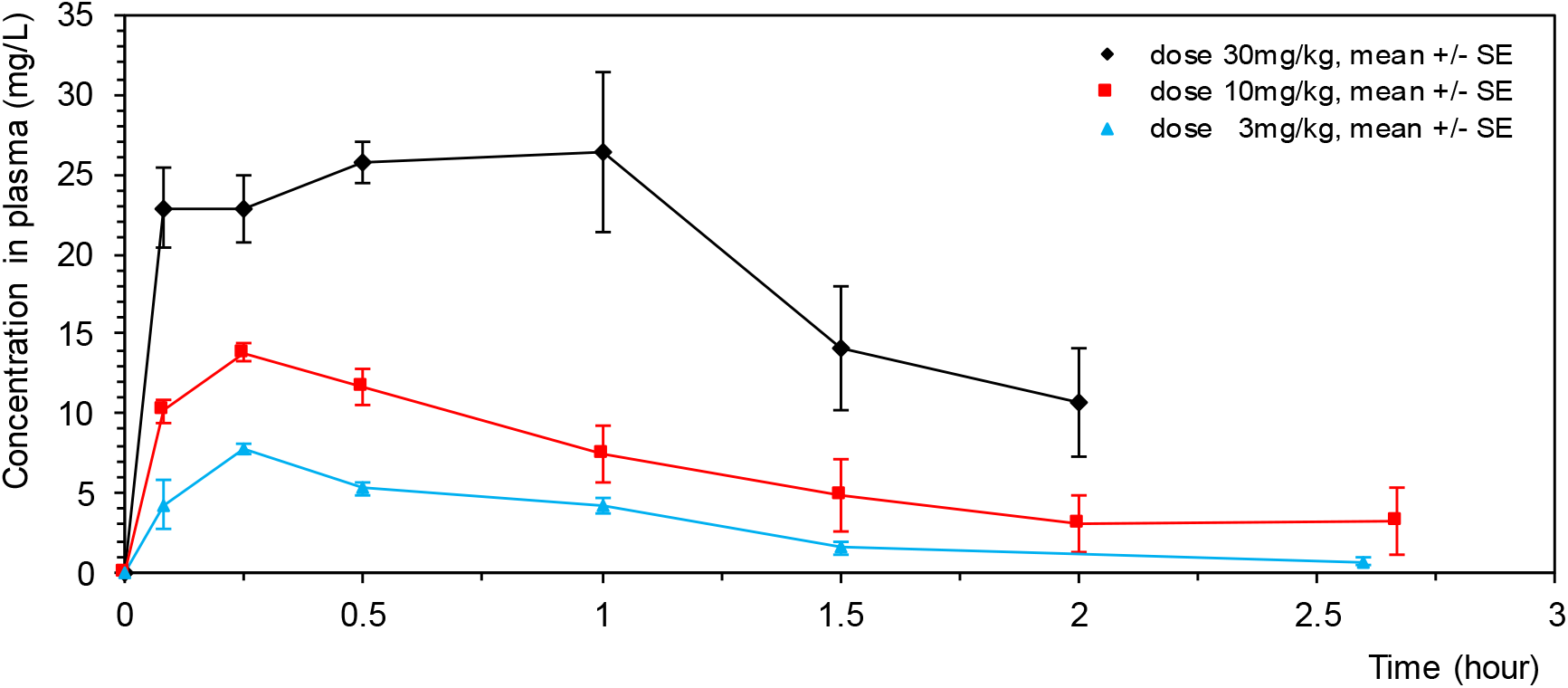
APC24-7 concentration in plasma after SC bolus dosage in thigh-infected female NMRI mice.

### Combination of Meropenem and APC24-7 is efficacious *in-vivo*

Combination therapy of APC24-7 and meropenem efficacy was tested in a neutropenic mouse thigh infection model against four strains -*E. coli* Espagne and AMA005000 and *K. pneumoniae* AMA2650 and ST258. The strains were selected based on demonstrating sensitivity to APC24-7 *in-vitro*, β-lactamase gene diversity, as well as demonstrating resistance to meropenem monotherapy *in-vivo* in pilot experiments (data not shown).

Initially, meropenem dose-response curves were established, with the aim of identifying sub-therapeutic meropenem monotherapy doses (data not shown). Combination therapy of sub-therapeutic meropenem doses with three dose levels of APC24-7: 3, 10 or 30 mg/kg, was then tested in an 8-hour infection model using two meropenem and APC24-7 administrations, at 2 and 6 hours after inoculation (Figure 6A-D). APC24-7 significantly improved meropenem efficacy in a dose-dependent manner against all four strains. Colistin 20 mg/kg QD or BID, estimated to provide clinically relevant exposure in the timeframe of the experiments [41], [42], was used as a positive control treatment.

**Figure 6.**
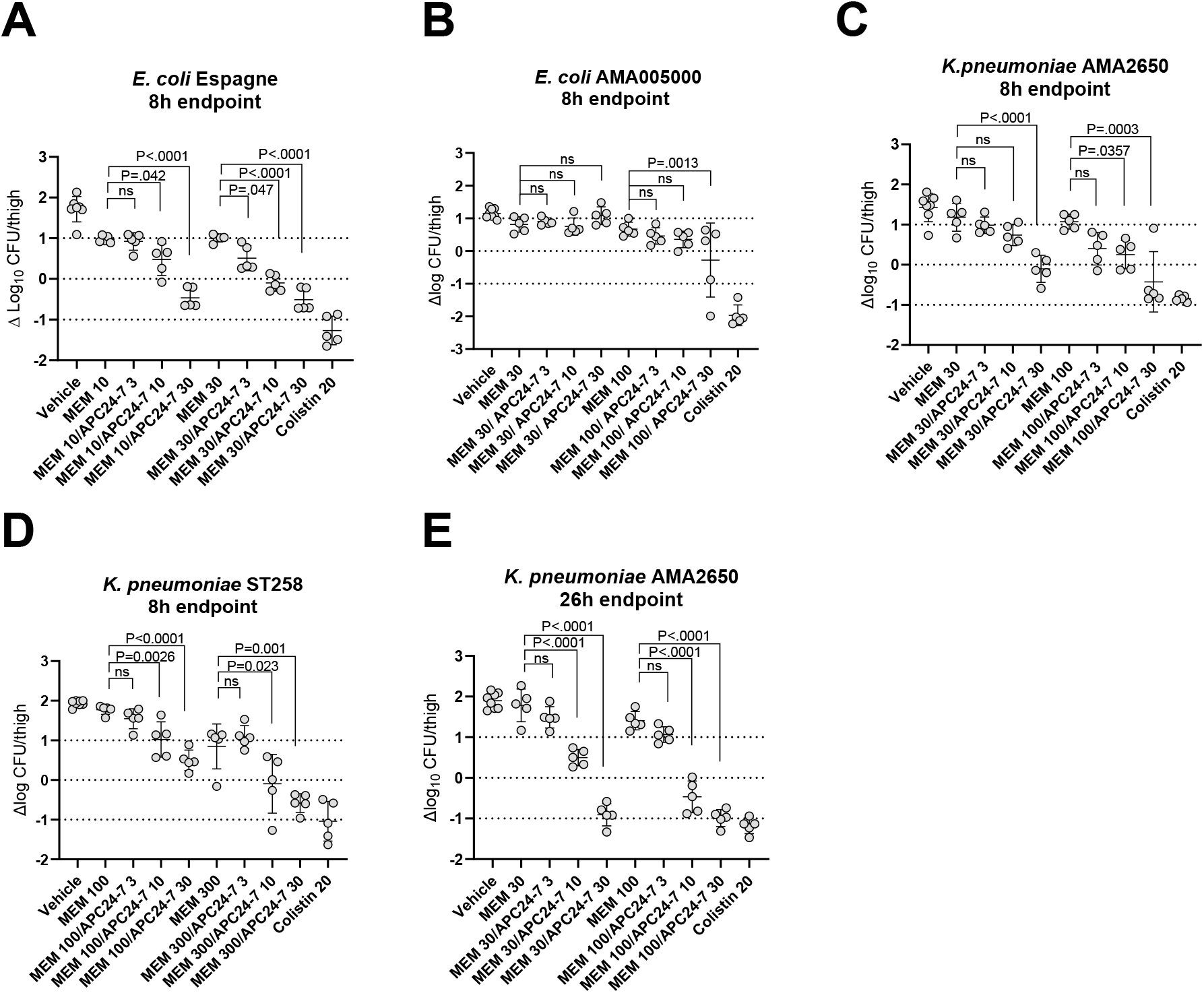
*In-vivo* effect of meropenem and APC24-7 combination treatment. Bacterial growth at 8 hours post- inoculation relative to the start-of-treatment baseline (approx. 6 log_10_ CFU for all isolates) is shown after two doses administered 2 and 6 hours after inoculation in mice infected with carbapenem-resistant isolates: (A) *E. coli* Espagne, (B) *E. coli* AMA005000, (C) *K. pneumoniae* AMA2650, and (D) *K. pneumoniae* ST258. (E) For *K. pneumoniae* AMA2650, an additional experiment assessed bacterial burden at a 26-hour endpoint following Q4h dosing initiated 2 hours after inoculation. Colistin was administered Q12h as a positive control. Statistical comparisons were performed against the corresponding meropenem monotherapy group using ANOVA with Sidak’s post-test. Individual doses are indicated in mg/kg (e.g MEM 100 = meropenem 100 mg/kg Q4h)

For isolate *E. coli* Espagne (VIM-1, MEM MIC=32 mg/L), 30 mg/kg APC24-7 reduced the mean bacterial burden by ≥ 1 log_10_ compared with meropenem 10 mg/kg monotherapy, reaching the start-of-treatment baseline (Figure 6A). Increasing the meropenem exposure with a 30 mg/kg dose, allowed a lower dose of 10 mg/kg APC24-7 to reduce the mean bacterial load to baseline (Figure 6A).

*E. coli* AMA005000 (NDM-5, OXA-48, CTX-M-3, TEM-1B, MEM MIC=128 mg/L) was less responsive to APC24-7 *in-vivo* with only the combination of meropenem 100 mg/kg and APC24-7 30 mg/kg reducing the mean bacterial burden to baseline (Figure 6B). *K. pneumoniae* isolates AMA2650 (KPC-3, SHV-62-like TEM-1-A-like, MEM MIC=128) and ST258 (KPC-2, OXA-9 like, SHV-5-like, TEM-1A-like, MEM MIC=64) were both highly responsive to APC24-7 with doses of 10 or 30 mg/kg reducing the mean bacterial load to below baseline depending on meropenem exposure (Figure 6C), and was as efficacious as colistin treatment.

A 26h infection model testing Q4h dosing of the combination therapy against *K. pneumoniae* AMA2650 confirmed the initial finding and demonstrated improved efficacy, reaching -1 log_10_ kill and ≥ 2 log reductions compared with meropenem monotherapy (Figure 6E).

## Discussion

β-Lactam antibiotics, particularly carbapenems such as meropenem, remain essential last-resort therapies for severe infections caused by Gram-negative bacteria. However, the growing prevalence of resistance mediated by β-lactamase production by Gram-negative bacteria poses a major threat to the clinical utility of carbapenems [11], [43]. The development of broad-spectrum β-lactamase inhibitors (*BLi*s), such as APC24-7 [38], represents a promising strategy to restore the activity of existing β-lactams by protecting them from degradation by diverse β-lactamase enzymes [44].

Furthermore, the emergence of resistance through mutations in clinically relevant SBLs such as KPC-3, SHV, and CTX-M-15 during treatment with available *BLis* [43], [45]–[48] highlights the ongoing adaptive pressure on current inhibitors and reinforces the urgent need for continuous development of novel BL/*BLi* adjuvants. APC24-7 was recently described as a hybrid broad-spectrum *BLi* that restores the antibiotic activity of meropenem against carbapenem resistant *E. coli* and *K. pneumoniae*. In particular, the authors described a superior activity towards NDM-9 and IMP-26 enzymes, when compared to taniborbactam but poor activity against OXA-48 [38]. Here we further explored the effectiveness of APC24-7 in restoring meropenem antimicrobial activity, both *in-vitro* and *in-vivo* against clinical *Enterobacteriaceae* strains. Our findings provide preclinical proof-of-concept supporting APC24-7 as a dual MBL/SBL inhibitor capable of restoring meropenem activity across multiple resistance backgrounds. In checkerboard MIC assays, APC24-7 restored meropenem activity against clinical isolates of *K. pneumoniae* and *E. coli* with high meropenem MICs ranging from 16 to >1024 mg/L and expressing a broad selection of beta lactamase enzymes. For most K. pneumoniae isolates with very high meropenem MICs (≥256 mg/L), while lowering the meropenem MIC several fold, APC24-7 did not lower the meropenem MIC to the susceptible breakpoint even at 64 mg/L and for three out of the 31 K. pneumoniae isolates (9.7%), no impact on meropenem MIC was observed at this APC24-7 concentration. While all three unresponsive isolates expressed OXA-48, other isolates expressing the same enzyme was sensitized by APC24-7 and further work is needed to understand how carbapenem resistant *K. pneumoniae* avoids APC24-7 inhibition.

The diminished kill rate over time as seen in Figure 4 panel A-B, suggests that the inhibition is not sustained and could point to *BLi* instability, induced enzyme expression or possibly the emergence of a resistant subpopulation, and will have to be investigated further. It is also important to note that zMIC values derived from the first 3 to 5 hours represent early-phase pharmacodynamics, when β-lactamase inhibition is maximal. For this reason, zMIC values should not be compared directly with the 16-20 h MIC values in Table 1 & 2, which inherently include regrowth and differ conceptually from zMIC. Differences between these metrics are therefore expected in systems where inhibitor activity decreases over time. Moreover, the initial net growth rate data indicate that increasing APC24-7 concentrations beyond 32 mg/L does not yield additional killing in *E. coli* IHMA997800. Taken together, these isolate-specific observations may help explain why, across the broader panel of strains, the impact of APC24-7 on meropenem activity was variable.

Non-enzymatic resistance mechanisms, such as efflux regulation, target modification, and signaling pathways involved in β-lactamase sensing [12], [49] may also be present and contribute to the observed resistance phenotype. In this context, and as part of an exploratory analysis to assess potential non-enzymatic contributors to carbapenem resistance, a subset of *K. pneumoniae* clinical isolates was screened for mutations in outer membrane porins, including ompK35, ompK36, and ompK37 [50]. Point mutations were frequently observed in ompK36 and ompK37, which may contribute to reduced carbapenem susceptibility, whereas ompK35 remained conserved (supplementary Table S1), however these results were not correlated with checkerboard results.

The *in-vivo* experiments reinforced APC24-7’s potential as a *BLi*. Mouse thigh infection efficacy studies confirmed that APC24-7 rendered four highly meropenem resistant clinical isolates susceptible to treatment with meropenem at doses where meropenem monotherapy did not impact bacterial burden. For this proof of concept, we selected isolates where APC24-7 reduced the meropenem MIC to breakpoint in *in-vitro* assays. Follow-up experiments should focus on restoration of meropenem susceptibility using human simulated meropenem dosing regiments [51] and include isolates with very high meropenem MICs and for which the susceptibility breakpoint was not achieved with even 32 or 64 mg/L APC24-7. For one of the four isolates (*E. coli* AMA005000, NDM-5, OXA-48), *in-vivo* combination treatment was less effective despite checkerboard and time-kill analysis indicated that this isolate was no less sensitive to APC24-7. This particular result may reflect the contribution of the OXA-48 enzyme, against which APC24-7 exhibits limited activity or to differences in the function of BL enzymes *in-vivo* as compared to *in-vitro*.

Initial PK evaluation of APC24-7 showed that exposure was not linear. The half-life was approximately 40–50 minutes, which is prolonged compared with meropenem, which has a half-life of <15 minutes in mice. Although further pharmacokinetic and pharmacodynamic studies are required to establish the optimal dosing regimen, these findings suggest that APC24-7 is suitable for combination therapy.

Our data suggest that the combination of meropenem and APC24-7 may represent a complementary β-lactamase inhibition strategy, particularly for resistance backgrounds dominated by KPC and selected MBLs where current therapeutic options remain limited.

## Materials and Methods

### Bacterial isolates and compounds/chemicals

A panel of clinical isolates covering as broad range of BL enzymes was obtained from the Statens Serum Institut (SSI), the CDC antimicrobial resistance collection or kindly provided by Professor Morten Sommer from Novo Nordisk Center for Biosustainability, Technical University of Denmark. ATCC25922, ATCCBAA1705 and NCTC13443 were included as quality control strains.

Meropenem for injection was purchased from STADA, whereas APC24-7 was synthetized by Kappa solutions AS, acquired by Balchem Corporation.

### Determination of minimal inhibitory concentration (MIC)

The minimal inhibitory concentration (MIC) of meropenem and APC24-7, was determined by the microbroth dilution method in accordance to the guidelines established by the Clinical and Laboratory Standards Institute (CLSI) [39].

In summary, 96-well plates were utilized for two-fold dilutions of meropenem or APC24-7 in MH-broth. These dilutions ranged from 1 to 1024 m/mL for meropenem and 1 to 64 mg/L for APC24-7. Each well was then inoculated with the standard inoculum size recommended by CLSI. As part of the procedure, appropriate ATCC strains were included to serve as quality control (QC) strains. *E. coli* ATCC 25922 was used as meropenem MIC control and not included in other assays.

### Checkerboards assays

Checkerboard testing was conducted in 96-well microtiter plates following the methodology described by Bellio et al (2021) [52], with a few modifications. Plates were arranged with decreasing concentrations of meropenem along the x-axis and APC24-7 along the y-axis. Both the antibiotic and the inhibitor were assessed within a concentration range of 0.06 to 64 mg/L. In each well, 50 µL of bacterial suspensions, at approximately 10^6 CFU/mL, were combined with the antibiotic and inhibitor. Plates were then incubated at ± 35ºC for 18 hours.

### Time-kill experiments

Nine clinical isolates—*E. coli* IHMA997800, *E. coli* Espagne, *E. coli* AMA005000, E. coli 149, *K. pneumoniae* ST258, *K. pneumoniae* B68-1, *K. pneumoniae* AMA4817, *K. pneumoniae* 1149 and *K. pneumoniae* AMA2650—expressing different β-lactamases, were chosen for time-kill assays. These assays were conducted in duplicate, adapting the protocol outlined in the Clinical and Laboratory Standards Institute (CLSI) document M26-A [40].

Briefly, 96-well plates were inoculated with approximately 10^5 CFU/mL bacterial suspension. Meropenem was tested at 0.25, 1, 4, and 16 mg/L and MIC concentration of each strain. These concentrations were tested alone and in combination with relevant APC24-7 concentrations as determined based on checkerboard results for each individual strain.

The plates were incubated at ± 35 º C with shaking for 24 hours and samples taken at time points 0, 1, 3, 5, 8 and 24 hours, for CFU (colony forming units) determination.

Time-kill curves were generated by plotting mean logarithmic colony counts (log_10_ CFU/mL) against time to compare the 24-hour killing effects of meropenem monotherapy and meropenem-APC24-7 combination.

### Fitting of an Emax type of bacterial growth rate model to time-kill data

Equation 1 was used to estimate the initial net growth rate, *μ*, based on the mean slope defined by the first, few time points of CFU data (from *t* = 0 to *t* = *Δt*) for each curve, i.e. for each tested concentration *c* of meropenem (for fixed concentration of APC24-7). Values of 3 and 5 hours were used for *Δt* in all *μ* estimates.

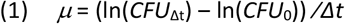

An Emax type of net growth rate model (Eqs. 2-3) was then fitted to the *μ* (*c*) data points:

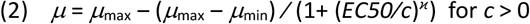

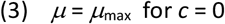

where *ϰ* is the negative slope of the sigmoid curve at meropenem concentration *c* = *EC50*.

In the parameter estimation procedure, *μ*_max_ was fixed to the value found for the growth control curve (*c* = 0 mg/L) which left three parameters free to be estimated: *μ*_min_, *EC50* and *ϰ*.

If *μ*_min_ < 0, a pharmacodynamic MIC, *zMIC*, can furthermore be defined as the concentration of the antibiotics at which *μ* = 0. The value of *zMIC* is closely related to EC50 and can be derived from Eq. 2. All parameter estimation was done with the TimKil program in the PKPDsim software package [53].

### Identification of point mutations in outer membrane protein genes

Whole-genome sequence assemblies of *K. pneumoniae* isolates were analyzed using ResFinder [54] to identify carbapenem resistance-associated point mutations in outer membrane protein–encoding genes. Default parameters were used.

### Bacterial inoculum for *in-vivo* studies

*K. Pneumoniae* and *E. coli* strains were propagated on 5% horse blood agar (SSI Diagnostica) overnight and the inoculum was prepared approximately one hour prior to inoculation by suspension of single colonies in sterile 0.9% saline to an optical density of 0.13 at 540 nm, giving a density of approx. 2 x 10^8 CFU/mL and then further diluted in 0.9% saline to 2 x 10^7 CFU/mL before injecting mice with 0.05 mL in the left thigh, i.e. approx. 10^6 CFU. For each experiment, the size of the inoculum was verified by making ten-fold dilutions in 0.9% saline, of which 20 µl was plated on 5% blood agar plates with subsequent counting of colonies after incubation overnight at ±35°C in ambient air.

### Mice and housing

Outbred female HsdWin:NMRI mice 26-30 gram (Inotiv, Netherlands) were housed at the animal facility at Statens Serum Institut. The temperature was 22°C +/-2 ^o^C and the humidity was 55 +/-10%. The mice were housed in individually ventilated type 3 macrolone cages with bedding from Tapvei. The air changes per hour were approximately 8-12 times (70-73 times per hours inside cages), and light/dark period was in 12-hours interval of 6 a.m.-6 p.m./6 p.m – 6 a.m. The mice had ad libitum access to domestic quality drinking water and food (Teklad Global diet 2916C-Envigo) and occasionally peanuts and sunflower seeds (Køge Korn A/S). Further, the animals were offered Enviro-Dri nesting material and cardboard houses (Bio-serv). Studies were ethically reviewed and adheres to the European Directive 2010/63/EEC and Danish license No. 2021-15-0201-01046.

### Pharmacokinetics study and bioanalysis

Pharmacokinetic studies in infected mice were carried out in female neutropenic NMRI 16 hours after an intramuscular infection as described below. APC24-7 was administered SC at 3, 10 or 30 mg/kg dose levels. Plasma was sampled at 5, 15, 30, 60, 120 minutes after dose and at approx. 160 min after dose where some mice met the humane endpoint due to the infection. In-life phase was at SSI and bioanalysis was done by Lablytica (Uppsala): A reversed-phase gradient high-performance liquid chromatography (LC) using a C18 analytical column with gradient elution (0.1% formic acid in water and acetonitrile) was used for separation. Detection was performed using mass spectrometry (MS) with positive ion spray ionization. An ion spray voltage of 1.0 kV, a cone gas of 30 L/h, and a desolvation gas of 1000 L/h were used on the MS. PK parameters were calculated by noncompartmental analysis using the initESTIM program in the PKPDsim software package [53].

### Murine thigh infection model

A previously described neutropenic thigh murine model was used for all experiments [31]. In brief, female NMRI mice were rendered neutropenic by cyclophosphamide pre-treatment and received an intramuscular injection with approx. 6 log_10_ CFU in the left thigh. Treatment was initiated 2 hours after inoculation and repeated q4h. Endpoint was either 8h (2 doses) or 26h (6 doses) after inoculation. Thighs were quantified for bacterial load at the 2h start-of treatment baseline in untreated animals and at endpoint following treatment. Bacterial isolates were selected based on ability to proliferate *in-vivo*, for their lack or low response to meropenem monotherapy as well as positive indication for APC24-7 and meropenem combination therapy in *in-vitro* experiments.

The mice were scored for clinical signs of infection throughout the study. In 8-hour, duration experiments, scores were mild across all groups. In 26-hour, duration experiments, scores were mild to moderate in vehicle controls, and no mice met the humane endpoint. The meropenem used was Stada powder for injection 500 mg. Meropenem and APC24-7 was admixed and given as a single injection. All injectables were kept refrigerated and protected from light.

### Statistical analysis

CFU data was log_10_ transformed prior to analysis. Samples that were below the level of quantification were assigned the LOQ value of 125 CFU/thigh i.e. 2.1 log_10_ CFU. ANOVA and Sidaks multiple comparison test were used to compare log_10_ CFU values for multiple groups. P-values < 0.05 were considered significant. All statistical comparison was made using the software GraphPad Prism 10 for Windows, GraphPad Software, CA USA.

## Abbreviations

BLs: β-lactam antibiotics
*BLi*: β-lactamase inhibitor
MBLs: metallo-β-lactamases
SBLs: serine-β-lactamases
CRE: carbapenem-resistant Enterobacterales

## Acknowledgements

We thank the technical assistance of technicians at Statens Serum Institute.

## Supplementary Figures / Tables

**Supplementary Figure 1.**
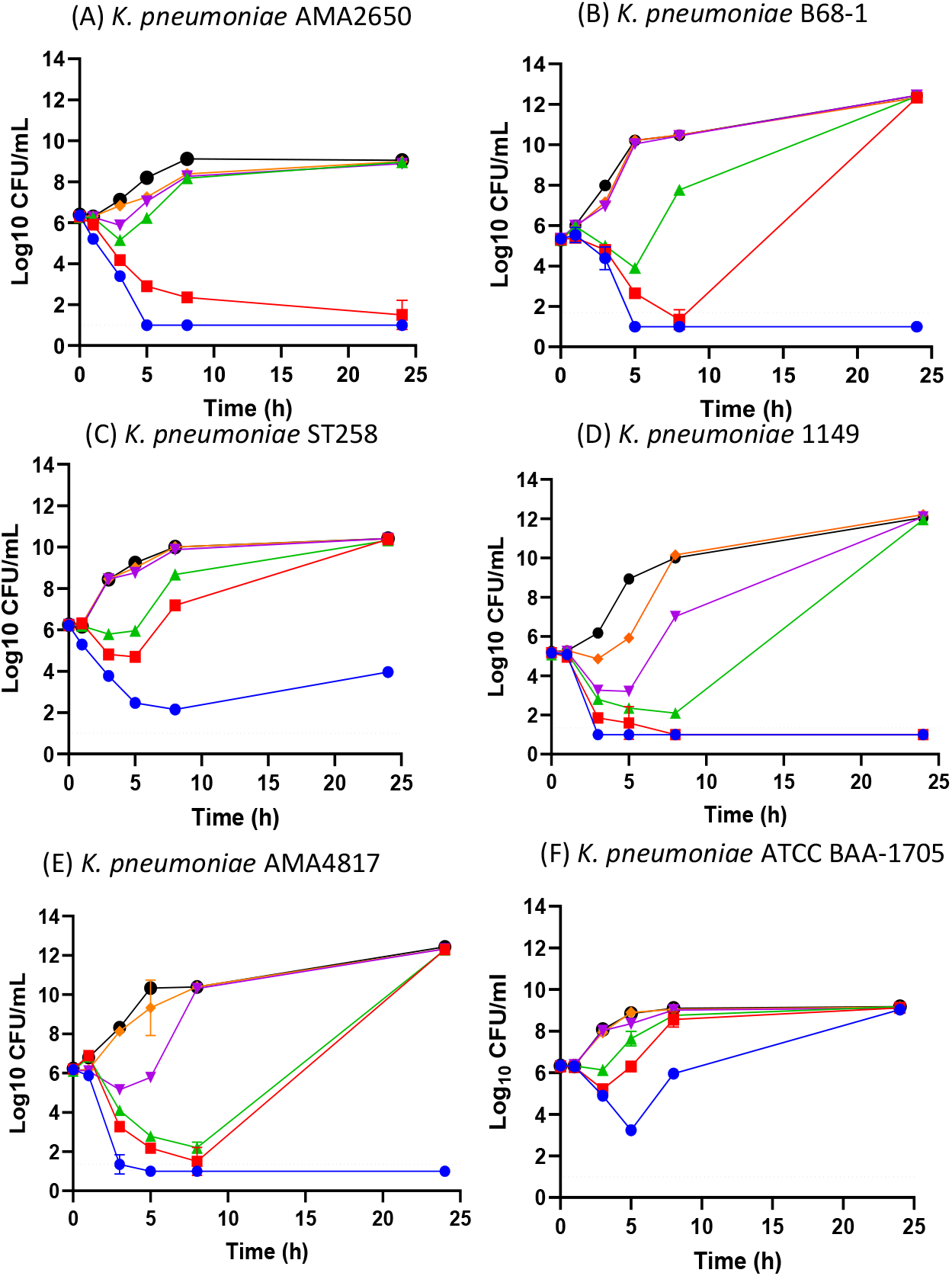
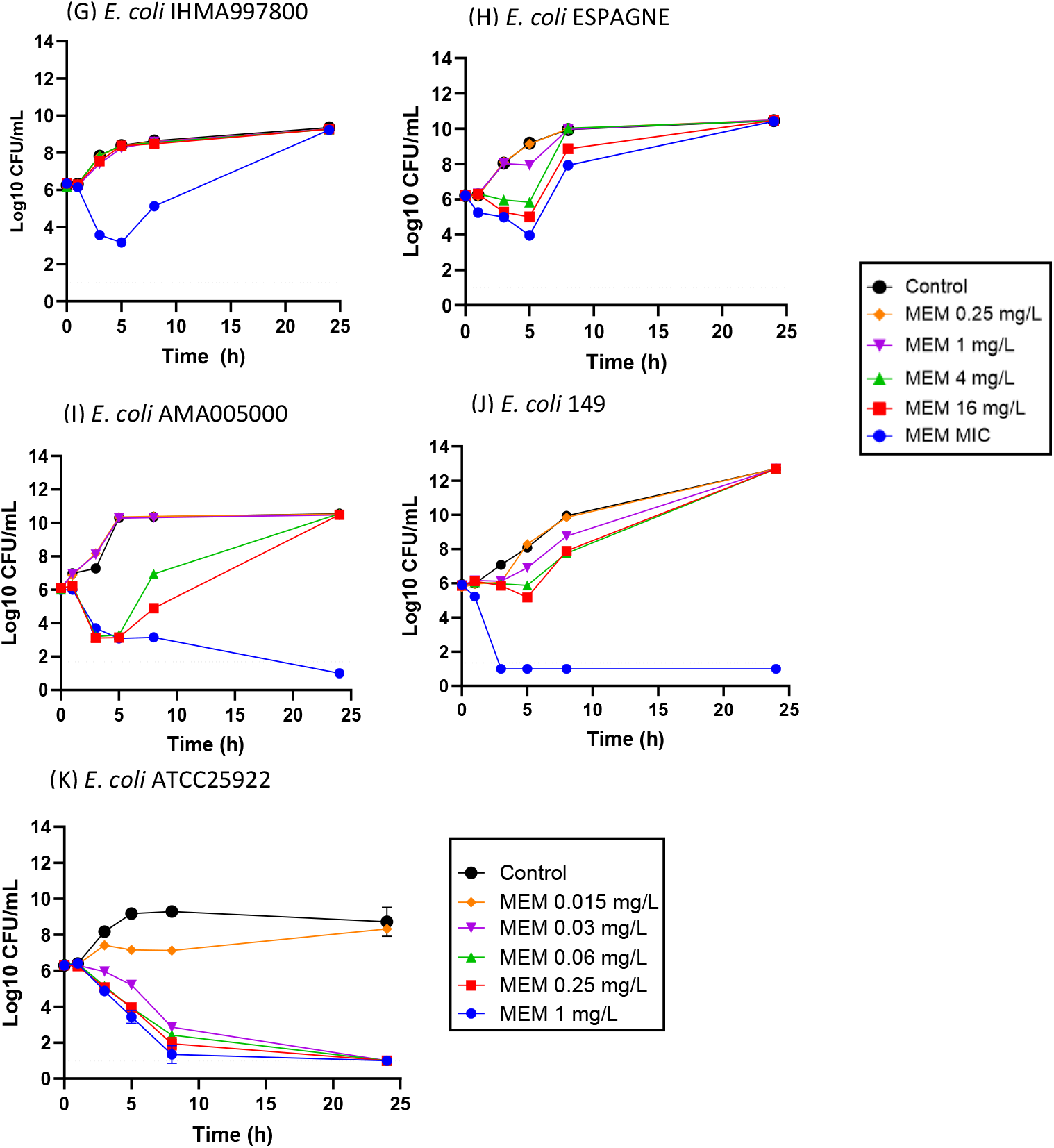
Time-kill curves of (A) *K. pneumoniae* AMA2650, (B) *K. pneumoniae* B68-1, (C) *K. pneumoniae* ST258, (D) *K. pneumoniae* 1149 and (E) *K. pneumoniae* AMA4817 (F) *E. coli* IHMA997800, (G) *E. coli Espagne*, (H) *E. coli* AMA005000, (I) *E. coli* 149 and (K) *E. coli* ATCC25922 (control), treated with meropenem alone at MIC value and 0.25, 1, 4 and 16 mg/L. Growth controls (drug free media) were included in all experiments. Dotted line on each graph indicates the lower detection limit. Data is shown as the average CFU/mL of duplicates per time point.

**Table S1.** Point mutations in outer membrane porins detected by RESFINDER.

| Isolates Id. | Point Mutations conferring resistance to carbapenems |
| --- | --- |
| *K. pneumoniae* ST258 | ompK37 (p.I70M, p.I128M, p.N230G) |
| *K. pneumoniae* DIH | ompK36 (p.N49S, p.L59V, p.L191S, p.F207W, p.A217S, p.N218H, p.D224E, p.L228V, p.E232R), ompK37 (p.I70M, p.I128M) |
| *K. pneumoniae* ST147 | ompK36 (p.A217S); omp37 (p.I70M, p.I128M) |
| *K. pneumoniae* AMA003821 | ompK36 (p.A217S); omp37 (p.I128M, p.N230G) |
| *K. pneumoniae* AMA1348 | ompK36 (p.A217S); ompK37 (p.I70M, p.I128M, p.N230G) |
| *K. pneumoniae* AMA2119 | ompK37 (p.I70M, p.I128M, p.N230G) |
| *K. pneumoniae* AMA877 | ompK36 (p.A217S); ompK37 (p.I70M, p.I128M, p.N230G) |
| *K. pneumoniae* AMA2650 | ompK36 (p.A217S, p.N218H); ompK37 (p.I70M, p.I128M) |
| *K. pneumoniae* AMA005047 | ompK36 (p.A217S), ompK37 (p.I70M, p.I128M) |
| *K. pneumoniae* AMA004435 | ompK36 (p.A217S), ompK37 (p.I70M, p.I128M, p.N230G) |
| *K. pneumoniae* CPO20170084 | ompK36 (p.A217S), (p.I70M, p.I128M) |
| *K. pneumoniae* AMA004208 | ompK36 (p.A217S, p.N218H); ompK37 (p.I70M, p.I128M) |
| *K. pneumoniae* AMA004950 | ompK36 (p.A217S), ompK37 (p.I70M, p.I128M, p.N230G) |
| *K. pneumoniae* AMA004443 | ompK36 (p.A217S); ompK37 (p.I70M, p.I128M, p.N230G) |
| *K. pneumoniae* AMA004417 | ompK37 (p.I70M, p.I128M) |
| *K. pneumoniae* AMA4817 | ompK37 (p.I70M, p.I128M, p.N230G) |
| *K. pneumoniae* NCTC13443 | ompK36 (p.A217S, p.N218H); ompK37 (p.I70M, p.I128M) |

